# EMERGENCE OF OXA-48-LIKE CARBAPENEMASES IN CLINICAL ENTEROBACTERALES FROM GUATEMALA

**DOI:** 10.1101/2024.04.04.588149

**Authors:** Carmen J. Mazariegos, Anna Quinto, Andres Herrera, Sara E. Barillas, Laura R. Valenzuela, Gabriela Ordóñez, Paola Sierra, Carolina Arevalo, Christine Seah, Roberto G. Melano

## Abstract

Five isolates (four *Klebsiella pneumoniae* and one *Escherichia coli*) displaying resistance to piperacillin/tazobactam and ertapenem and reduced susceptibility to imipenem were found positive for *bla*_OXA-48_-like genes (*bla*_OXA-181_ and *bla*_OXA-232_) carried on plasmids described previously. Although two *K. pneumoniae* recovered in 2020 were clonally related, the results suggest that *bla*_OXA-48_-like genes have been circulating in Guatemala since 2019. This is the first report of this type of carbapenemase in Central America.

---

The oxacillinases or OXA ß-lactamases (Ambler class D) are a diverse group of enzymes with high hydrolytic activity towards semisynthetic penicillins such as oxacillin, hence the name OXAs. Some OXA-variants display weak carbapenem hydrolysis, being the OXA-48-like family the most commonly found in *Enterobacterales* (1). Within the OXA-48 group, the variants OXA-181 and OXA-232 are the second and third most abundant. OXA-232 was derived from OXA-181 with a single amino acid substitution, while the latter differs from OXA-48 by four substitutions, although it is not derived from it (2,3). OXA-48-like enzymes confer resistance to penicillins, narrow-spectrum cephalosporins, limited activity against extended-spectrum cephalosporins and most ß-lactam inhibitors (e.g., clavulanate, sulbactam, and tazobactam), and weak hydrolysis of carbapenems (4). Because of these features, their detection is a diagnostic challenge due to the range of antibiotic resistance patterns conferred to *Enterobacterales*, which are often underreported because they are considered as susceptible to some carbapenems according to international guideline cutoffs from the Clinical and Laboratory Standards Institute (CLSI) and the European Committee on Antimicrobial Susceptibility Testing (EUCAST). However, in combination with other bacterial resistance mechanisms such as impermeability and/or production of extended-spectrum or AmpC β-lactamases, they can develop high resistance to carbapenems (4). In addition, some OXA-48 variants show improved activity against broad-spectrum cephalosporins that lack carbapenemase capacity due to an amino acid deletion (e.g., OXA-163 and OXA-405), while others have improved their hydrolytic profile against carbapenems (e.g., OXA-162 and OXA-181) (4,5).

OXA-48 and its variants are common in certain regions of the world (e.g., the Middle East, North Africa, and European countries such as Belgium and Spain) (6), but they are relatively rare in Latin America (6-9). In Guatemala, the most common carbapenemase reported in *Enterobacterales* has been NDM and KPC (10,11). In the present study we describe the first reported cases of OXA-48-like producers isolated in the country and in Central America.

In 2019 and 2020, numerous Enterobacterales isolates were referred to the National Health Laboratory of Guatemala whose antibiotic susceptibility, analyzed with Vitek® 2 Compact (bioMérieux, Marcy-l’Étoile, France) and disc diffusion method, showed resistance to at least one carbapenem, according to CLSI guidelines (12). The ones displaying resistance to piperacillin/tazobactam and ertapenem and reduced susceptibility to imipenem were tested by molecular method. GeneXpert® Carba-R (Cepheid, Sunnyvale, CA) analysis revealed that five isolates (four *Klebsiella pneumoniae* and one *Escherichia coli*) were positive for *bla*_OXA-48_-like gene (Table 1). Using short- and long-read sequencing (Illumina and Oxford Nanopore, respectively), it was possible to sequence the entire genome and circularize plasmids and chromosomes, type the isolates, and identify all the AMR determinants as described (13).

**TABLE 1.**
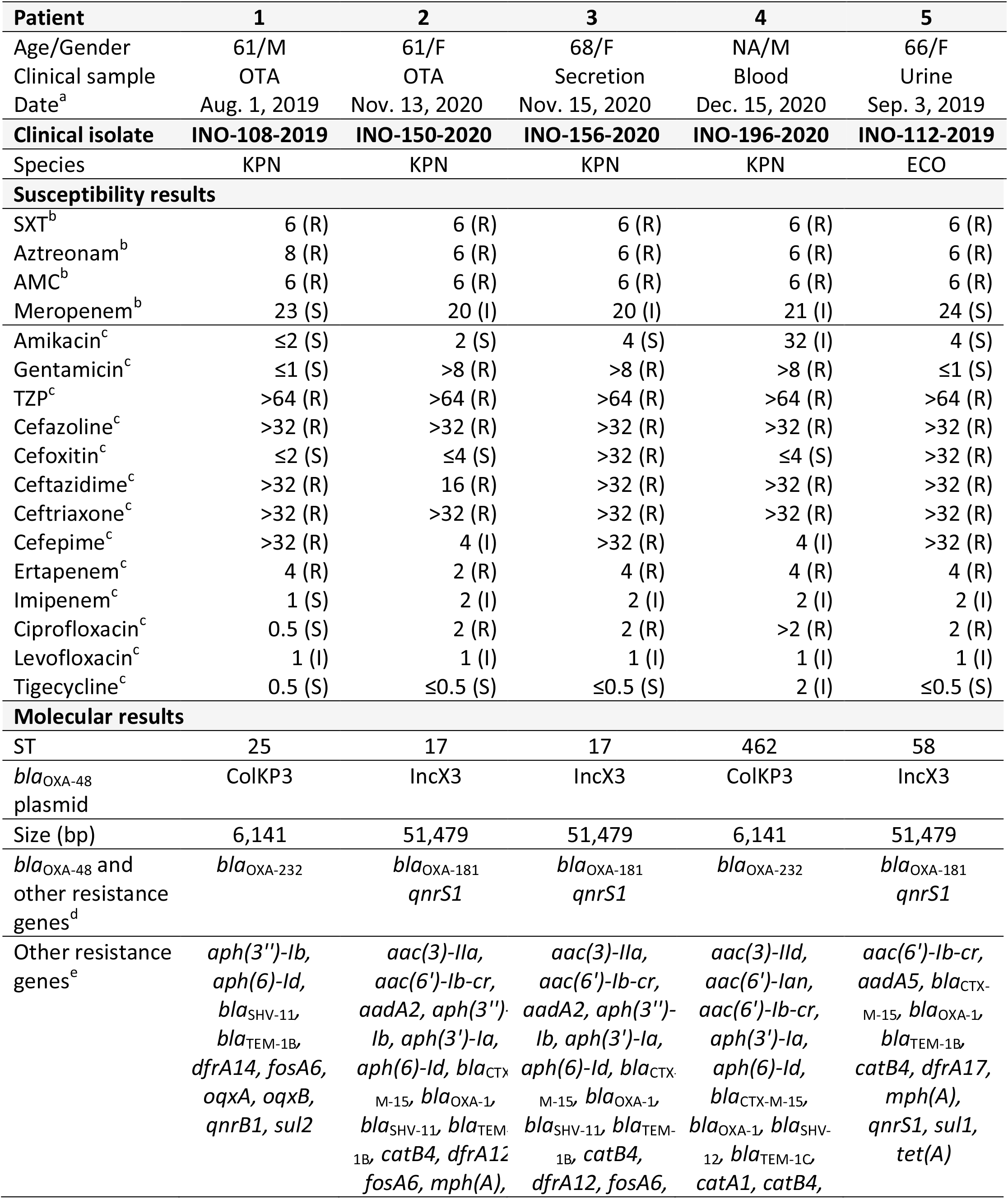
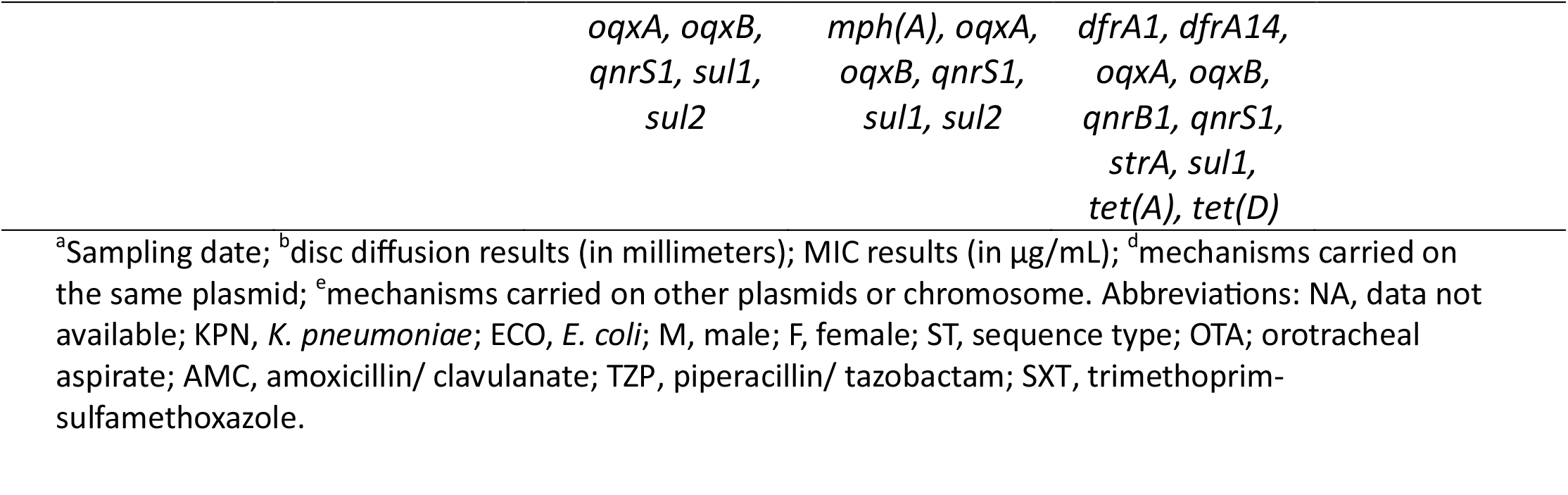
Data on OXA-48-like-producing isolates from Guatemala.

The antibiograms showed that all isolates were resistant to most the antibiotics tested, with the only exception of tigecycline and amikacin (four isolates susceptible and one intermediate) and levofloxacin (all the isolates with intermediate resistance) (Table 1). Regarding carbapenems, ertapenem was the clearest marker for OXA-48-like suspicion, since all the isolates were resistant to this ß-lactam (meropenem and imipenem were intermediate or susceptible). The mechanisms of AMR identified by WGS of these isolates were consistent with the described MDR phenotypes found (Table 1).

By multilocus sequence typing (MLST) analysis, three sequence types (ST) were identified for *K. pneumoniae* (ST17, n=2; ST25 and ST462), while the *E. coli* isolate belonged to ST58 (Table 1). By single nucleotide polymorphisms (SNP) analysis, both ST17 *K. pneumoniae* isolates (INO-150-2020 and INO-156-2020) were almost identical (only one single SNP of difference) (Figure 1). These isolates were recovered from 2 COVID-19 patients hospitalized at the same time in the adult intensive care unit, supporting the clonal dissemination of this strain between them. These isolates were not genetically and epidemiologically related to the other ST462 (INO-108 - 2019) and ST25 (INO-196-2020) *K. pneumoniae* isolates (Figure 1). *K. pneumoniae* ST17 has been associated with opportunistic hospital-acquired infections (14). The other two STs were occasionally found (14-17). *E. coli* ST58 has emerged as a globally disseminated uropathogen that belongs to the environmental/ commensal phylogroup B1 (18).

**FIGURE 1.**
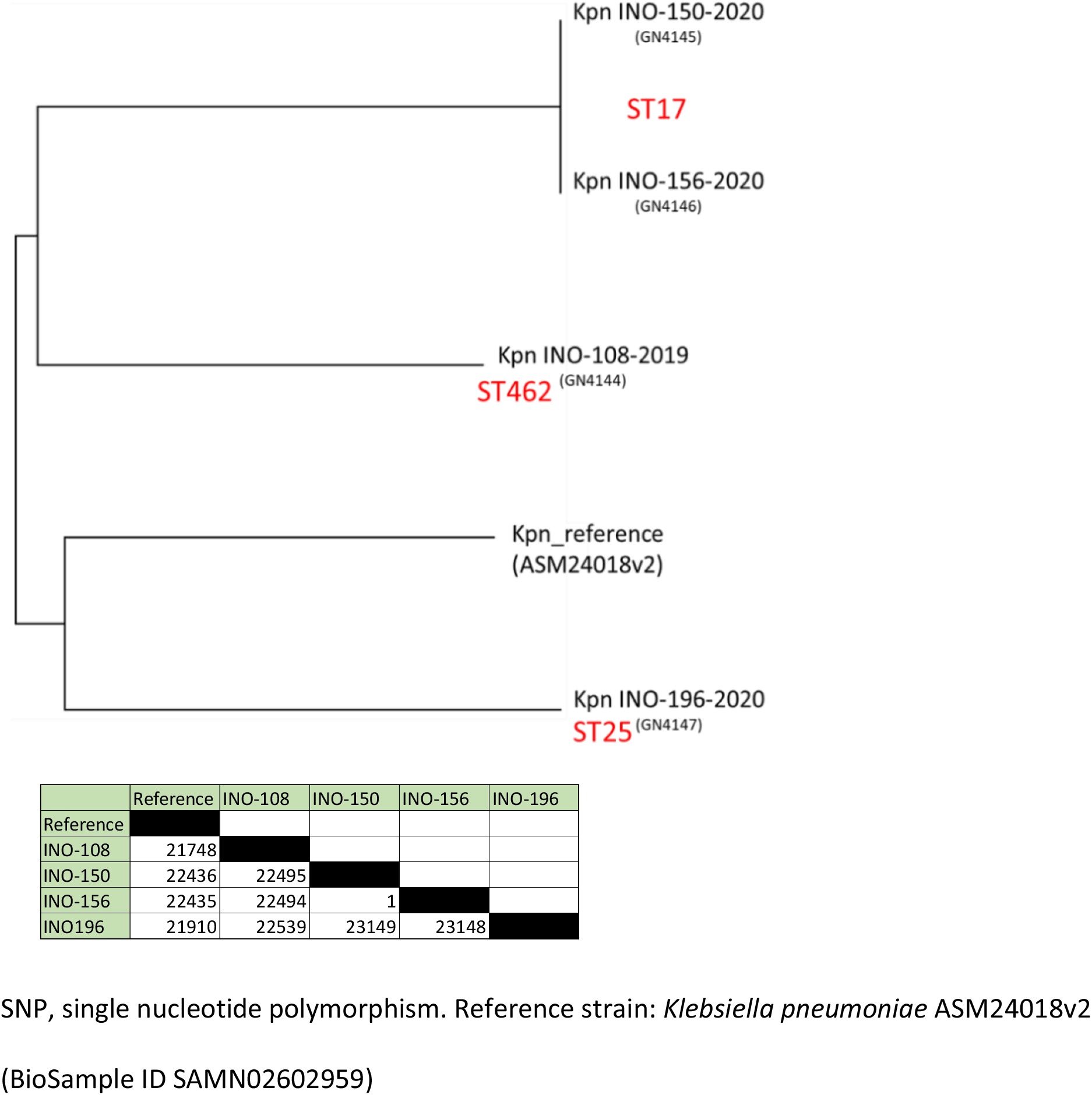
Phylogenetic tree based on OXA-48-like SNPs from Guatemala, and phylogenetic distance matrix according to the number of SNPs

IncX3 plasmids of 51,479 bp harboring *bla*_OXA-181_ were detected in the *E. coli* and both ST17 *K. pneumoniae* isolates (100% nucleotide identity). *E. coli* was not epidemiologically linked to the *K. pneumoniae* isolates, thus excluding horizontal dissemination. These IncX3 plasmids shared >99% nucleotide identity with other IncX3 plasmids in the GenBank (BioProject number PRJDB5126; GenBank accession no. MG570092 and KP400525.1) (19-21). The other two *K. pneumoniae* (ST25 and ST462) harbored identical ColKP3 plasmid of 6,141 bp carrying *bla*_OXA-232_ gene, with no nucleotide differences with previously described plasmids (BioProject no. PRJNA221868; GenBank accession no. JX423831) (3,22).

The first two patients (1 and 5, table 1) had no previous travel history, they carried different species of Enterobacterales, and variants of *bla*_OXA-48_ (*bla*_OXA-181_ and *bla*_OXA-232_) located in very different plasmids. In addition, the bacterial isolates were obtained from two different institutions located in distant cities. These data strongly suggest the independent acquisition of the first cases, which would be already circulating in the country since 2019. Due to the lack of information on the epidemiology of OXA-48-like producers in Central America, the probable origin of these isolates is unknown, being the first report of this type of carbapenemase in the subregion. Laboratories must be vigilant due to the difficulty of detecting the weak carbapenemase activity of OXA-48-like ß-lactamases. While synergy testing and colorimetric assays offer potential solutions, the variable nature of these enzymes can lead to false negative results. Accurate identification strategies are critical as under-detection affects treatment outcomes and complicates our understanding of global epidemiology.

## Data availability

The sequence data for the five isolates described in this study are available at NCBI under BioProject number PRJNA1062784.

